## supplement text and figures for "Evaluation of information flows in the RAS-MAPK system using transfer entropy measurements"

**Supplementary text**

**TE and conditional TE**

Statistical entropy:

The uncertainty of the system variable $x$ in a multistate system X can be described by the statistical entropy H(X), which is calculated from the probability distribution of $x$, p($x$) as follows:

$$H(X) = \sum_x -p(x)\log p(x).$$

Since uncertainty disappears once a measurement specifies the value of $x$, statistical entropy refers to the amount of uncertainty reduction, i.e, the amount of information obtained by the measurement.

Mutual information:

When there is a relationship (correlation) between two coexisting systems, X, Y (with system values $x$ and $y$), identifying the state of one reduces the uncertainty about the state of the other. This reduction is called mutual information, MI(X;Y). The mutual entropy can be calculated from p($x$), p($y$), and the joint distribution of $x$ and $y$, p($x$,$y$):

$$MI = H(X) - H(X|Y) = H(X) + H(Y) - H(X,Y) = \sum_x \sum_y p(x,y) log \frac{p(x,y)}{P(x)p(y)}.$$

H(X|Y) is the entropy of X conditioned on Y (see later). MI is a symmetric value with respect to X and Y, and means the amount of information that X (or Y) has with respect to Y (or X).

Transfer entropy:

Assuming that X and Y are time-varying, we denote their values at time $t$ as $x(t)$ and $y(t)$ and consider estimating $y(t)$ from $y$ at a past time Tr, $y$(Tr). Tr may not be a single time point but may have a certain time width. As above, the estimation accuracy can be described as the mutual information MI($Y_t$; $Y_{Tr}$). For this estimation, we need a simultaneous distribution of $y$(Tr) and $y(t)$.

If we have a tripartite simultaneous distribution of $x$(Tc), $y$(Tr), and $y(t)$, we can calculate the mutual information, MI($Y_t$; $X_{Tc}$, $Y_{Tr}$), using the distribution of $y(t)$ and the joint distribution of $x$(Tc) and $y$(Tr). Here we estimate $y(t)$ using information from $x$(Tc) and $y$(Tr) simultaneously. The difference between MI($Y_t$; $X_{Tc}$, $Y_{Tr}$) and MI($Y_t$; $Y_{Tr}$) is called the transfer entropy, TE, from X to Y.

$$TE_{X\to Y} = MI(Y_t; X_{Tc}, Y_{Tr}) - MI(Y_t; Y_{Tr}) = MI(Y_t; X_{Tc}|Y_{Tr})$$

Due to the non-negativity of the mutual information, TE is non-negative. Significantly large $TE_{X\to Y}$ means that the accuracy of the estimation of $y(t)$ has increased by considering the information in $x$(Tc). When Tc, Tr < $t$, a significantly large $TE_{X\to Y}$ indicates that X(Tc) has some sort of causality to Y($t$). TE is asymmetric for X and Y as shown in the definition above. $TE_{X\to Y}$ is a quantity that expresses how much X(Tc) controls Y($t$), independently of Y(Tr) (control capability). Note, however, that TE may appear due to spurious correlations. Even in this case, the TE still provides some insight into the system.

Conditional TE:

If another system, Z is measured at the same time as X and Y, we can consider the distribution of $x$ (or $y$) and the joint distribution of $x$ and $y$ under a given value of Z. These are the conditional probability distributions given z, p($x$|$z$) and p($x,y$|$z$). The mutual information calculated from the conditional probability distributions is called conditional mutual information.

$$MI(X;Y|Z) = \sum_z \sum_x \sum_y p(z)p(x,y|z)log\frac{p(x,y|z)}{P(x|z)p(y|z)}$$

We can calculate conditional TE from conditional mutual information:

$$TE_{X \to Y|Z} = MI(Y_t; X_{Tc}, Y_{Tr}|Z) - MI(Y_t; Y_{Tr}|Z).$$

By the definition of MI:

$$TE_{X \to Y} = H(Y_t|Y_{Tr}) - H(Y_t|X_{Tc}, Y_{Tr}), \text{ and}$$

$$TE_{X \to Y|z} = H(Y_t|Y_{Tr}, Z) - H(Y_t|X_{Tc}, Y_{Tr}, Z).$$

We defined the difference TE (ΔTE) as:

$$TE_{X \to Y|z} - TE_{X \to Y} = \{H(Y_t|Y_{Tr}, Z) - H(Y_t|Y_{Tr})\} + \{H(Y_t|X_{Tc}, Y_{Tr}) - H(Y_t|X_{Tc}, Y_{Tr}, Z)\}.$$

In general, due to the non-negativity of the information carried by Z, the uncertainty in $Y_t$ does not change or decrease after conditioning on Z, i.e, $H(Y_t|W,Z) \leq H(Y_t|W)$. Here, W is any kind of random variable. Therefore, the value in the first parenthesis of ΔTE is non-positive and the value in the second parenthesis is non-negative. A negative ΔTE means that at least the first value is negative, i.e., Z correlates with $Y_t$ independently of $Y_{Tr}$. While a positive ΔTE means that at least the second value is positive, i.e., Z correlates with $Y_t$ independently of $X_{Tc}$ and $Y_{Tr}$. A tandem reaction path, $Z \to X_{Tc} \to Y_t$ does not satisfy the latter condition, so its ΔTE cannot be positive. But ΔTE can be negative because Z has a control path to $Y_t$ that is not mediated by $Y_{Tr}$. Among the simple networks of X, Y, and Z, the pathways where Z controls $Y_t$ independently of $X_{Tr}$ and $Y_{Tc}$ to have the potential to produce positive ΔTE in the forward direction are $X_{Tc} \leftarrow Z \to Y_t$ and $X_{Tc} \to Z \to Y_t$ (Fig. 3E).

TE analysis in biology

Here are some examples of TE analysis applied to biology. TE analysis can be applied to any multidimensional data, including the results of molecular dynamics simulations. Kamberaj and van der Vaart (2009) used TE analysis to extract the causality of correlated motions within single molecules from molecular dynamics simulations of a DNA-binding protein molecule, ETS-1 transcription factor. They found that the direction of information flow changes upon DNA binding. This work revealed the mechanism of allosteric protein dynamics.

Wibral et al. (2011) analyzed TE in MEG data from a human auditory short-term memory experiment to estimate the network topology of brain modules. The topology was task type specific. Stetter et al. (2012) developed a method to infer the neural network connectivity from the TE between the series of single-cell calcium responses in the network. This method outperformed previous methods in simulations, and was applied to the experimental data of cultured neuronal cell populations. Ursino et al. (2020) developed another TE-based method for estimating network structure and connection strengths that is applicable to complicated 2-4 neuron networks. They reported that in the liner regime of the cell response, there was a fairly good correlation between TE and synaptic strength when the data lengths were sufficient. These are examples of the application of TE to neural circuit analysis.

Pahle et al.(2008) performed stochastic simulations of the association between $Ca^{2+}$ binding to a protein and developed a framework for investigating the relationship between the patterns of $Ca^{2+}$ dynamics and the ability to transfer information from the $Ca^{2+}$ dynamics to the $Ca^{2+}$ binding level of the protein. Ito and Sagawa (2015) showed theoretically that the robustness of an information processing system to environmental changes can be quantitatively evaluated by TE. They analyzed the experimentally obtained adaptive behavior of E. coli chemotaxis signal transduction, and found that chemotaxis signal transduction is efficient as an information transmission device despite being highly dissipative as a thermodynamic engine.

**Stochastic ODE simulations**

Models and simulation:

We consider reaction networks between X, Y, and Z (Fig. S11A):

$$\frac{dY}{dt} = (a_{12}X(t) - a_{32}Z + a_2)(T_Y - Y) - b_2Y + \xi_Y,$$

$$\frac{dZ}{dt} = (a_{13}X(t) - a_{23}Y + a_3)(T_Z - Z) - b_3Z + \xi_Z.$$

Here, X is the input to Y and Z set to be a single-peaked two-phase function as follows:

$$\frac{dX_0}{dt} = -a_0X_0 + \xi_{X0},$$

$$\frac{dX_1}{dt} = (a_0X_0 - a_{21}Y - a_{31}Z) - b_{01}X_1 + \xi_{X1},$$

$$\frac{dX_2}{dt} = (a_0X_0 - a_{21}Y - a_{31}Z) - b_{02}X_2 + \xi_{X2},$$

$$X(t) = X_1(t) + X_2(t)/5$$

For numerical simulations of the above model, $a_0$, $b_{01}$, and $b_{02}$ were set to 1, 0.1, and 0.005, respectively, to mimic the phosphorylation dynamics of EGFR in real cells qualitatively (Fig. S10). $a_2 = a_3 = 0.01, b_2 = 0.5, b_3 = 0.1,$ and $T_Y = T_Z = 10$. $\xi_i$ is a random parameter obeying a Gaussian distribution with the average = 0 and the standard deviation = $D\sqrt{F(t)}$ for each *dF/dt*, and *D* is a variable parameter controlling the noise level. *D* = 0.01 was used for all simulations. Other parameter values are listed in Fig. S11A. By setting some parameter values to zero, we changed the network topology of the models. All of the models we examined here contain an input path from X to Y. $X_i$ shown in Fig. S11A is the initial value of $X_0$ and controls the input strength. These parameter values were chosen arbitrarily to obtain transient responses of Y and Z with unimodal intensity distributions in the range of $X_i$.

For each model, four simulations of reaction time courses were performed by varying $X_i$ from 0.02 to 0.35 in increments of 0.05 (Fig. S11B). In the simulations, Y and Z were first equilibrated in the absence of input (X(t) = 0) for $t$ = -50 to 0 (5,000 steps in the simulations) from appropriate initial values, and then Y(0) and Z(0) were used for the initial values of the simulation with input X($t$ > 0). The trajectories of Y and Z were calculated 500 times for each $X_i$ during $t$ = -3~60.

TE and conditional TE:

First, TE time courses between Y and Z were calculated for each model using all data sets varying $X_i$ as the averages of 100 bootstrap runs (Fig. S11B). In the tandem reaction model (X→Y→Z), significant amounts of TE were observed in the forward (Y→Z) direction, as expected. Amounts of bw-TE (Z→Y) were small and should be a statistical fluctuation. Adding a feedback loop from Z to Y or X (feedback models: X→Y↔Z and X→Y→Z→ X) increased bw-TE amounts. The size of TE in the parallel input model $(Y \leftarrow X \rightarrow Z)$ and depended on the reaction rate constants. When the reaction rate constant from X to Y ($a_{12}$) was larger than that from X to Z ($a_{13}$), fw-TE was dominant, but when $a_{12}$ was smaller than $a_{13}$, bw-TE became dominant. These results are not surprising given the spurious causality between Y and Z in this model. The X-mediated model $(Y \leftrightarrow X \rightarrow Z)$ does not contain feedback loops from $Z$ to $Y$, and its bw-TE level was low. Thus, simulations suggest that detection of a significant amount of bw-TE requires a feedback loop from Z or parallel inputs to Y and Z from X.

Next, the difference between TEs (ΔTE) conditioned and non-conditioned to $X_i$ was examined (Fig. S11C). Conditional TE is the average of the TEs for each $X_i$ value, while non-conditional TE is the TE for mixed data for all $X_i$ values. We performed 100 times bootstrapping for the non-conditional and conditional TEs and calculated ΔTE = conditional TE – non-conditional TE for each bootstrap run. ΔTE was calculated only between the statistically significant TE values. Significant positive ΔTE was observed only in the forward direction of the Z to X feedback model (X→Y→Z→X), where input X mediates the information path from Y to Z as expected (Y→Z→X→Y→Z). Negative significant ΔTE has been observed in some conditions where X is connected to $Z_t$ (in fw direction) or $Y_t$ (in bw direction) with skipping $Z_{Tr}$ or $Y_{Tr}$, respectively. This is also in line with expectations. However, it is not easy to explain what determines the sign and amount of ΔTE.

### Supplemental Figures

**Figure S1. Calculation of transfer entropy.**

The procedure for the calculation of TE from SOS(Tc) to RAF(*t*) with reference to RAF(Tr) is shown. (Here, Tr = Tc.) From the single-cell dynamics of SOS and RAF translocations (left), the single-cell response distributions **SOS**(Tc), **RAF**(Tr), and **RAF**(*t*) were extracted (middle to upper right; cell number = n). Then, the covariance matrix between the extracted distributions (bottom right) was calucted, from which the TE was calculated according to the formula shown in the 'Materials and Methods' section. R: RAF, S: SOS, c: TC, r: Tr. ($\mathbf{X}_i$, $\mathbf{Y}_j$): covariance beween **X**(i) and **Y**(j). ($\mathbf{X}_i$, $\mathbf{X}_i$) is the variance of **X**(i).

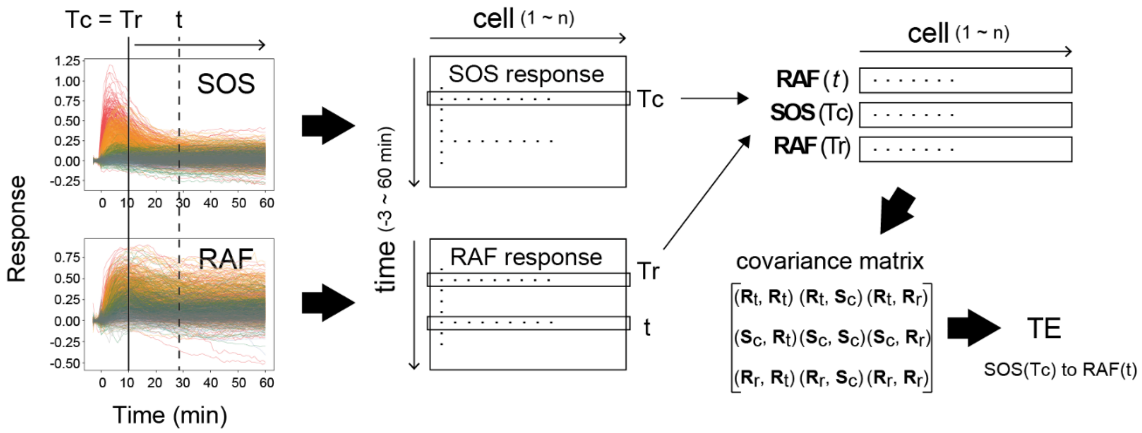

**Figure S2. Measurement reproducibility**

**A.** Single-cell response intensities of SOS and RAF at three different time points after EGF stimulation, with EGF dose indicated. Different colors represent measurements obtained on different days. Responses on each day overlap with a small bias. **B**. Numbers of cells shown in A. **C.** Inter-day variation in response mean. Variation was defined as the absolute value of the difference between the daily mean and the overall mean divided by the 5-95% range of the total single-cell response. The dashed line indicates a 30% variation.

A

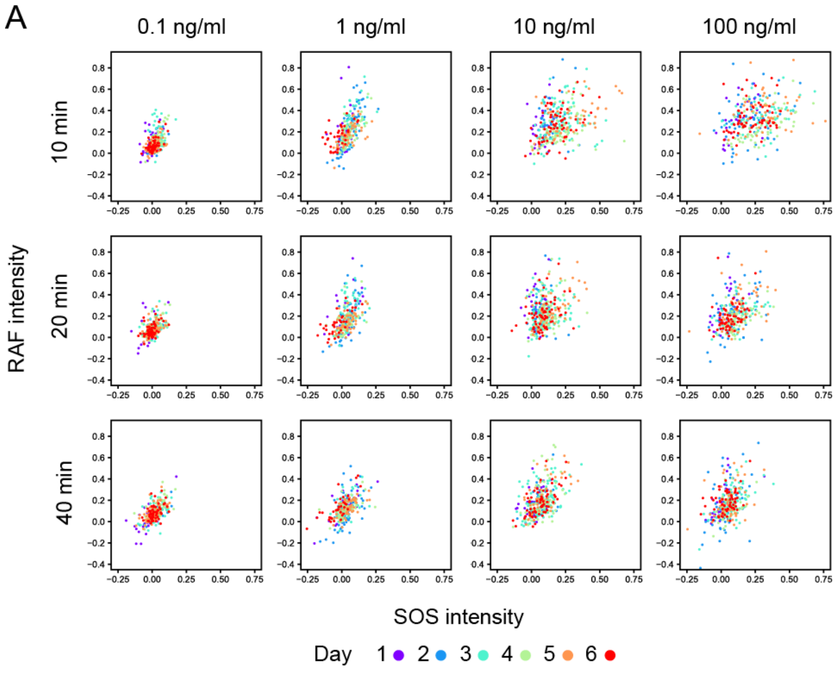


B

Number of cells

| EGF | day1 | day2 | day3 | day4 | day5 | day6 |
|---|---|---|---|---|---|---|
| 0.1 ng/ml | 27 | 22 | 77 | 63 | 59 | 52 |
| 1 ng/ml | 37 | 18 | 61 | 53 | 77 | 64 |
| 10 ng/ml | 31 | 37 | 105 | 75 | 52 | 70 |
| 100 ng/ml | 30 | 67 | 62 | 61 | 51 | 66 |

C

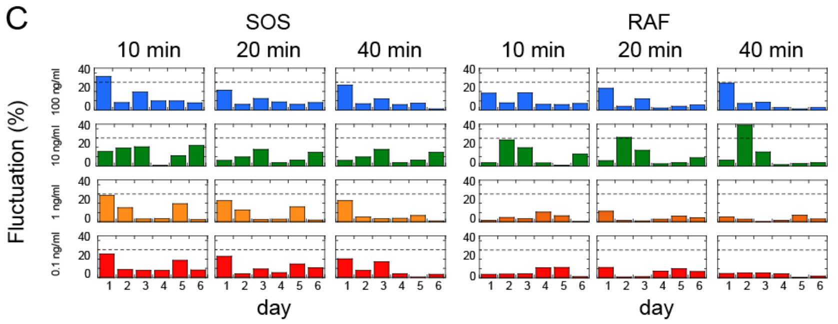

**Figure S3. Reactions and TEs under vehicle stimulation.**

**A.** Time courses of SOS and RAF responses after vehicle stimulation. These are the same plots shown as the black lines in Figure 1C. The upper panels show single-cell trajectories (n = 150). The bottom panels show the mean with SD. No significant responses were observed. **B.** Time courses of TE as the average of 100 times bootstrapping. The TE values are one order of magnitude lower than those observed after EGF stimulation (Fig. 2). These values could be noise caused by the positive bias of TE or true information transfer between basal responses in quiescent cells. Since there were no obvious temporal structures, it is less likely that the TEs were caused by the effect of the addition of vehicle solution.

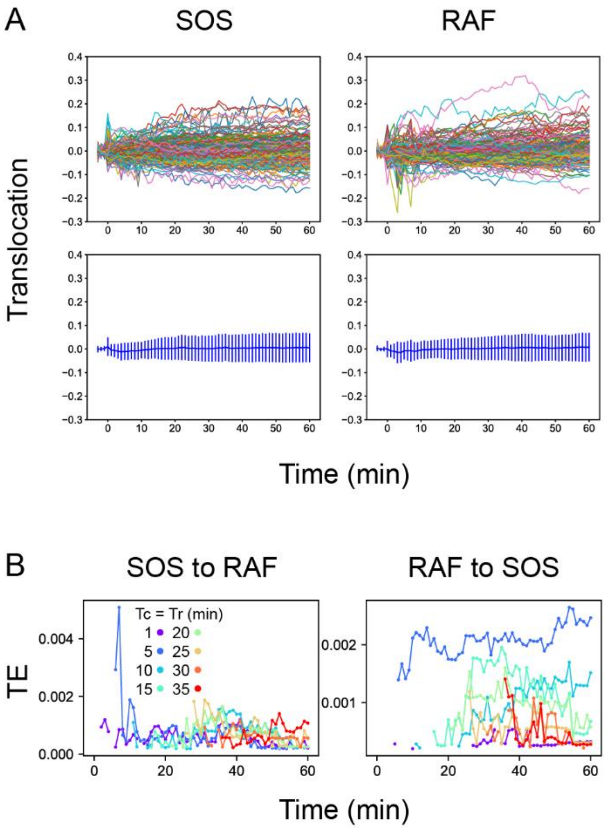

**Fig. S4. 3D response intensity distributions.**

Single-cell intensity distributions at the representative Tc, Tr, and *t* in TE time courses are shown as the 2D-projections. Colors indicate the densities of single-cell data points in a normalized unit.

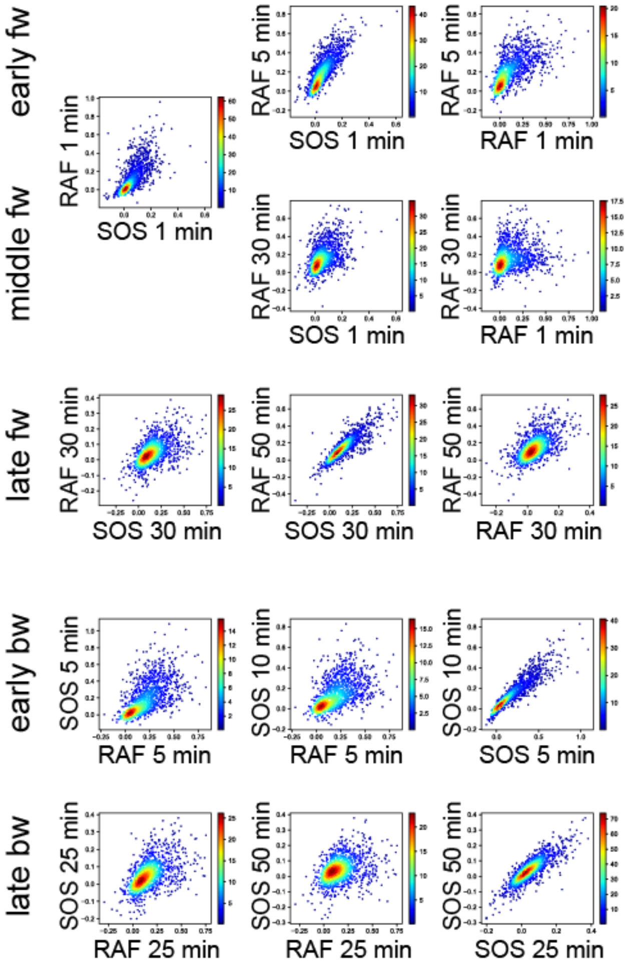

**Figure S5. Mutual information**

**A.** Time courses of the mutual information between the reaction intensities were calculated using the total 1317 data sets (Fig. 1D). The mutual information (MI) contents, the values of the two terms of TE (Fig. 2A) at the times Tc, Tr, and *t* used for TE calculation (Fig. 2B) are shown. Values higher than the upper 1% values in the 1000 times bootstrapping calculation under the null hypothesis were plotted. (In this criterion, all values were statistically significant in these calculations.) **B.** The significance threshold for the calculation in **A**, which is an indication of the error values in **A**. The thresholds for MI(X(Tr); X(t)) (~0.001) were significantly smaller than the values of TE(Y(Tc) to X(t)) (~0.01), supporting that the results of TE calculations were meaningful and sufficiently larger than the calculation errors in the subtractions between the two terms of TE.

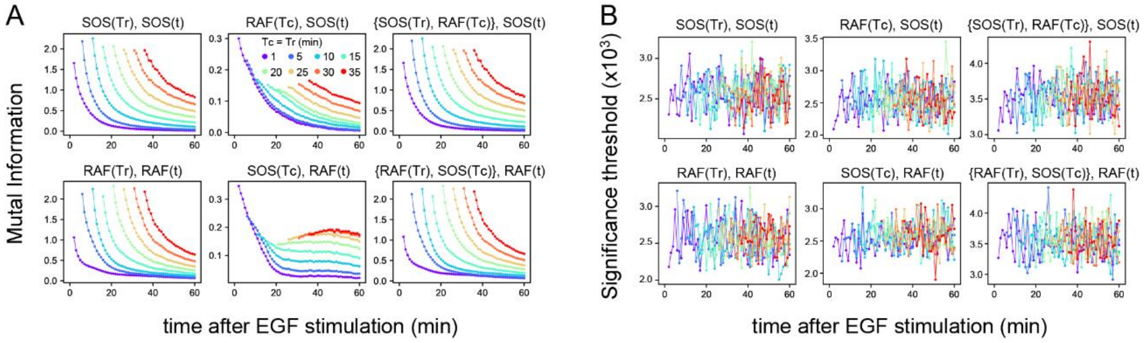

**Figure S6. Effects of the data number on transfer entropy detection**

Since the TE takes only non-negative values and the TE time courses are noisy, we checked if the limited data number caused a false detection of significant TE, reducing the number of data sequences used for calculation. Here, 100 to 800 data sets were randomly selected from the total 1317 data sets (Fig. 1D). **A.** Examples of TE time courses calculated using different numbers of data sets (n). As the number of data sets decreases, the number of time points where a statistically significant level of TE was detected also decreases. (These plots only show the statistically significant values.) This result means that TE was not detected falsely due to the limited number of data. **B.** TEsum values for 100 times of bootstrapping as a function of the data number. Values are normalized to those for 1317 data sets. The average (solid line) and 5 and 95% values (dotted lines) are shown. Even though the decrease in the data number caused the increase in the uncertainty (5–95% range), the averages were still within the 83–104% range of the TE for the total of 1317 data sets (excluding those for the early fw peak), when n ≥ 150. For the early fw peak, the bootstrap averages were within the 63–107% range.

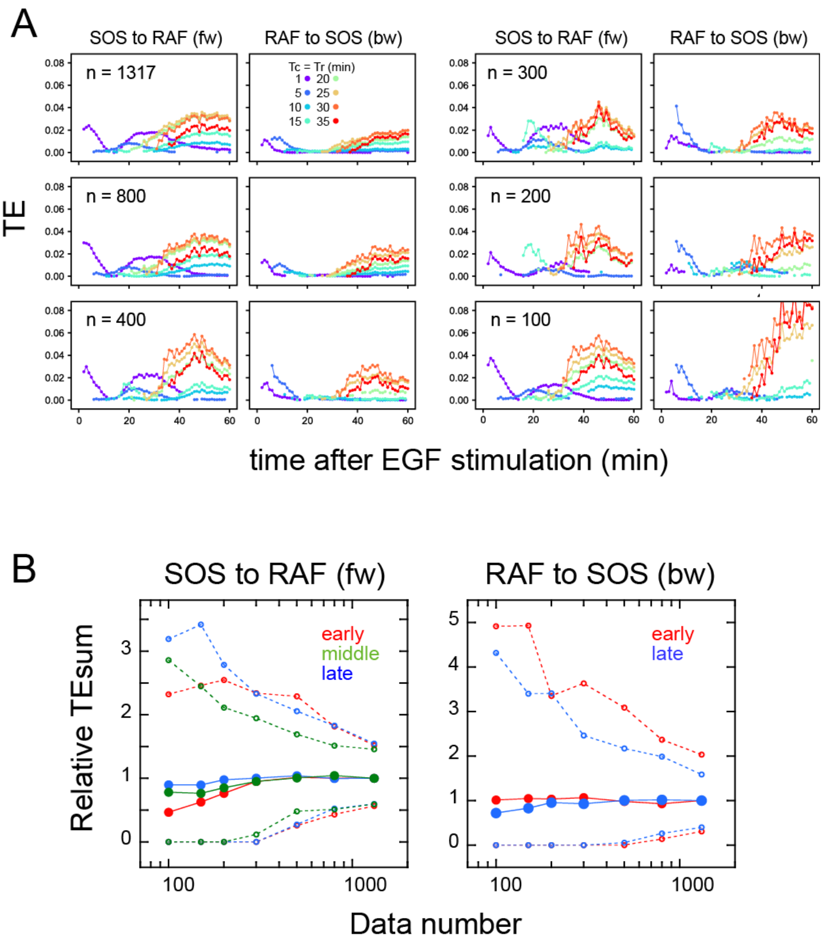

**Figure S7. TE exchange between SOS and RAF**

**A.** Time courses of TE were calculated for the SOS and RAF response using the total 1317 reaction dynamics data obtained under 0.1–100 ng/ml EGF (Fig. 1D). The calculation started from Tc = 1, and the time indicated by the arrows in the obtained TE time courses was used for Tc in the next round of calculation in the reverse direction. **B.** Diagram of TE exchange between SOS and RAF. Periods of the typical TE peaks are indicated.

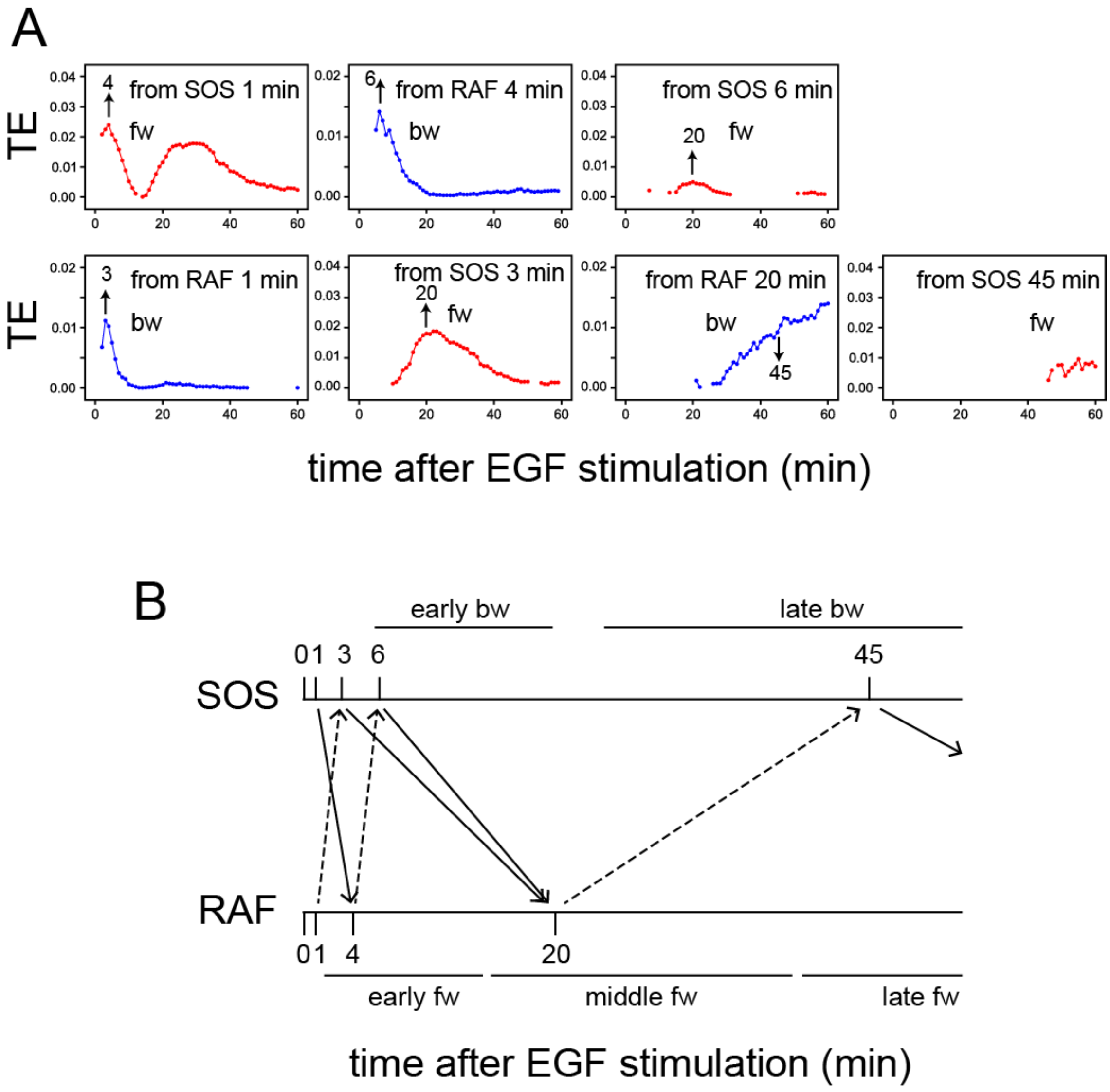

**Figure S8. EGF dose dependency in the TE time courses**

TE time courses were calculated for the SOS and RAF response time series at the indicated concentrations of EGF. n: data number. The calculations were performed for the time interval Tc = Tr from 1 to 35 min after the cell stimulation. Only statistically significant TE values (as defined in the Materials and Methods section) are plotted.

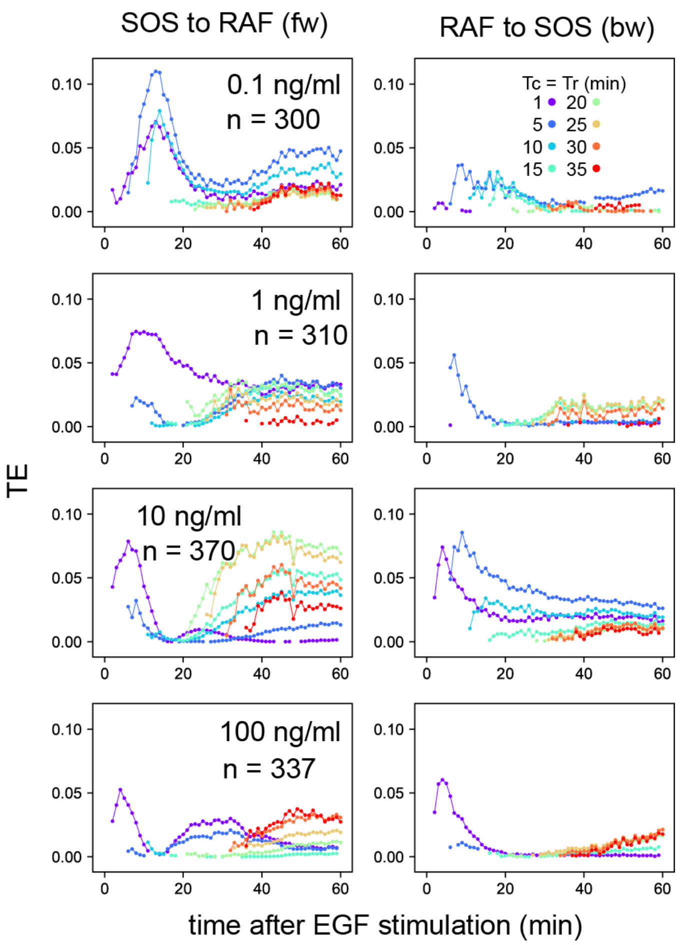

**Figure S9. Effects of the MEK inhibitor in cells with a Noonan syndrome SOS**

**A.** Single-cell time courses of SOS and RAF responses for Figure 5B. **B.** TE time courses for the data shown in (**A**). Calculations were performed for the indicated Tc = Tr from 1 to 35 minutes after cell stimulation. Only statistically significant TE values are plotted. (See Materials and Methods.) **C.** Difference between TEsum values in the presence and absence of the MEK inhibitor in R1131K cells. Positive values mean larger TEsum in the presence of the MEK inhibitor. Averages for 100 times bootstrapping are shown with the 5–95% interval. **bootstrapping $p < 0.05$. **D.** A possible pathway carrying middle fw-TE through a feedback loop. Action points of the MEK inhibitor (i) and R1131K mutation (K) are indicated.

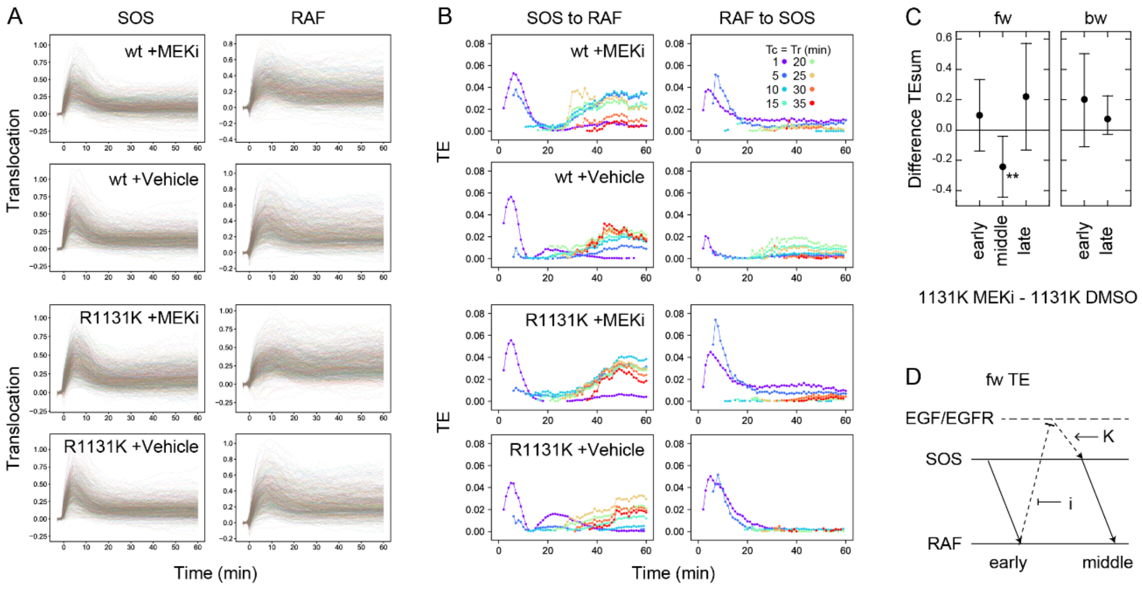

**Figure S10. Time course of EGFR phosphorylation**

Phosphorylation levels of Y1068 in EGFR indicating EGFR activity were measured. Representative Western blotting results (**A**) and quantification of four independent experiments (**B**) are shown. Cells expressing wt SOS and GFP-RAF were prepared under the same conditions as for SOS and RAF translocation measurements and stimulated with EGF. EGF dose and time after EGF stimulation are shown. Staining intensities were normalized to those for actin in the same sample and further to those of the same control sample of pEGFR (indicated by C). Phosphorylation levels peaked at 2-10 min after the EGF stimulation and decreased thereafter as expected from SOS, RAF response, and TE analysis.

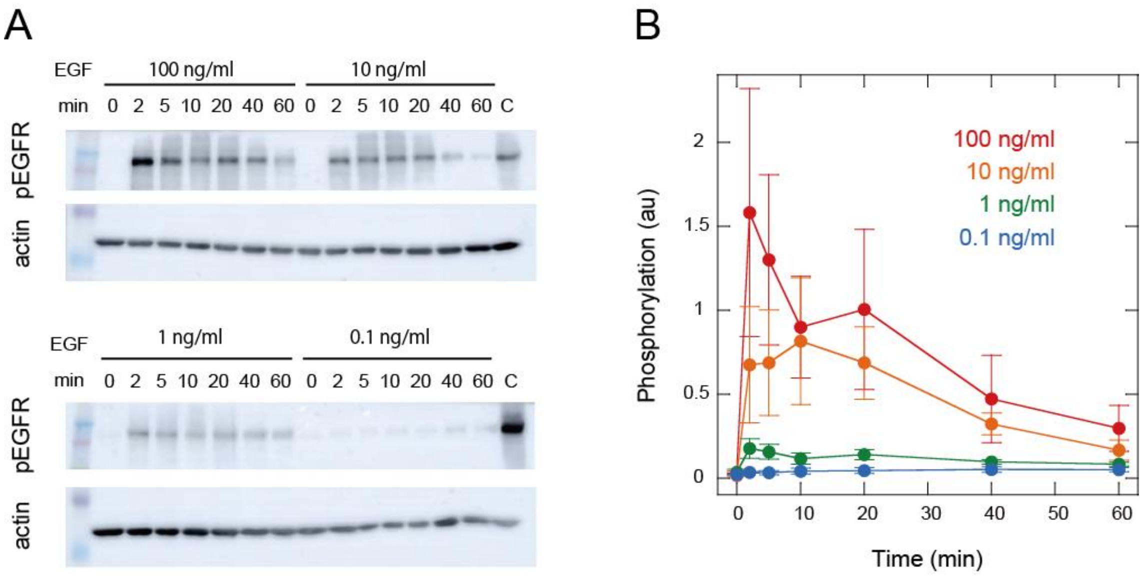

**Figure S11. Stochastic simulation and TE calculation for simple reaction networks**

**A.** Network structure and reaction parameters. $X_i$ is the initial intensity of $X_0$ and controls the amplitude of X (see Supplement text). **B.** Response time courses of X and Y. Averages of 500 simulated trajectories are shown with SD for each $X_i$ value. **C.** TE time courses for the mixed data under all $X_i$ values. **D.** Differences between conditional and non-conditional TE to $X_i$. **C, D.** The averages of 100 times bootstrap runs are shown. Only statistically significant values are plotted.

A

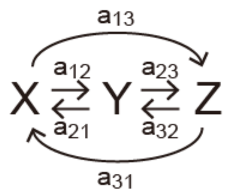


| Model | $a_{12}$ | $a_{21}$ | $a_{13}$ | $a_{31}$ | $a_{23}$ | $a_{32}$ |
|---|---|---|---|---|---|---|
| X→Y→Z | 0.5 | 0 | 0 | 0 | 0.1 | 0 |
| X→Y⇄Z | 0.5 | 0 | 0 | 0 | 0.1 | -0.03 |
| X→Y→Z (Z→X) | 0.5 | 0 | 0 | $-0.03X_i$ | 0.1 | 0 |
| Y←X→Z (a12 > a13) | 0.5 | 0 | 0.1 | 0 | 0 | 0 |
| Y←X→Z (a12 < a13) | 0.1 | 0 | 0.5 | 0 | 0 | 0 |
| Y⇄X→Z | 0.5 | $-0.03X_i$ | 0.1 | 0 | 0 | 0 |

B

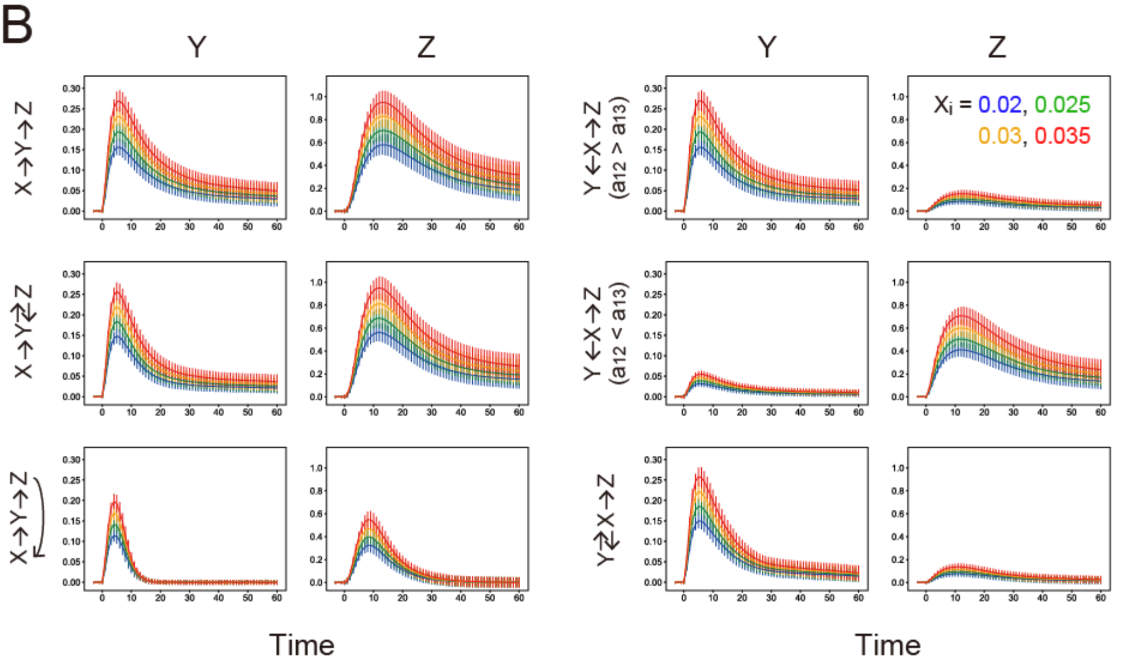

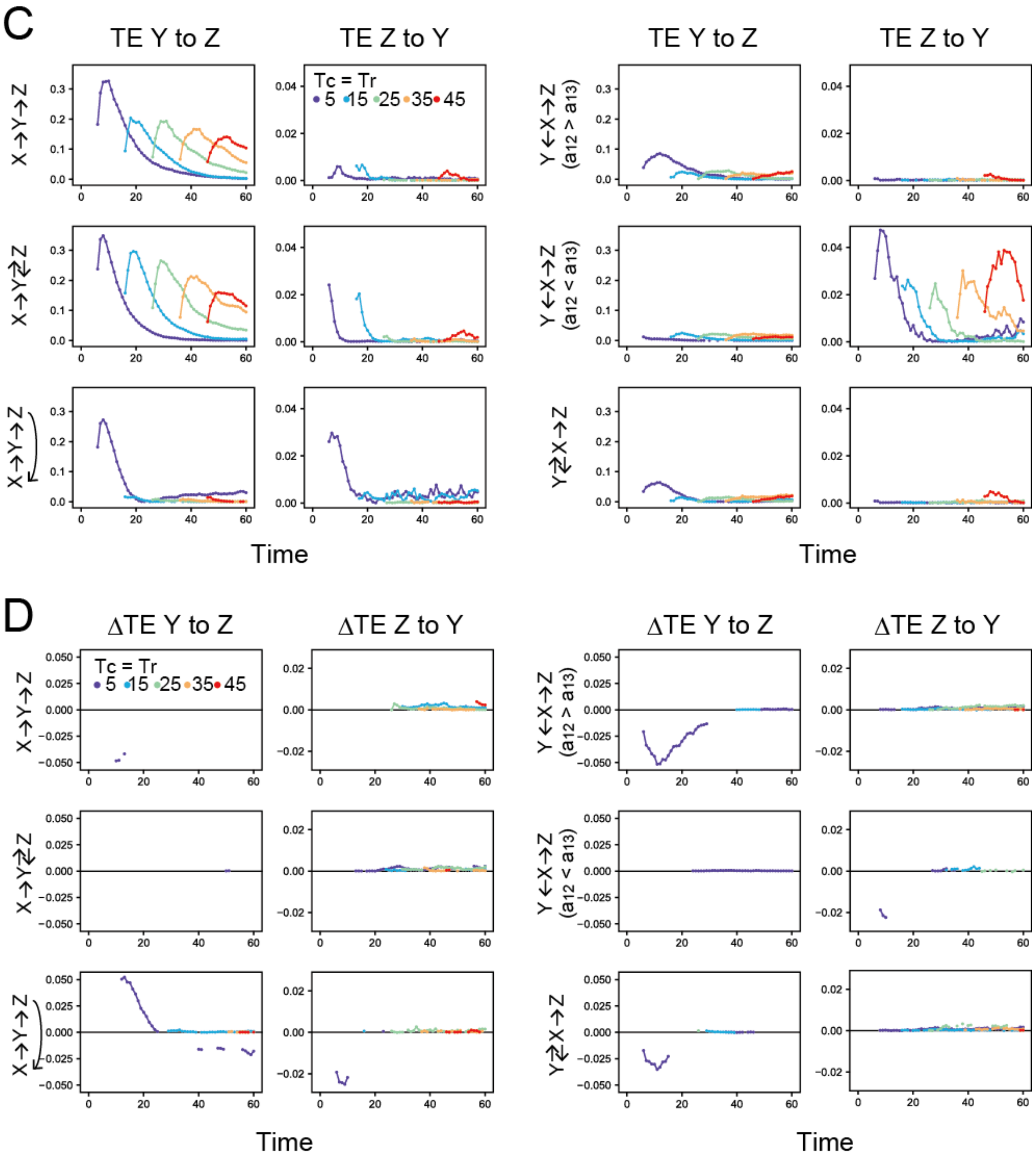
C
TE Y to Z
TE Z to Y
TE Y to Z
TE Z to Y
Tc = Tr
5 15 25 35 45
X→Y→Z
X→Y⇄Z
X→Y→Z
Y←X→Z
(a12 > a13)
Y←X→Z
(a12 < a13)
Y⇄X→Z
Time
Time
D
ΔTE Y to Z
ΔTE Z to Y
ΔTE Y to Z
ΔTE Z to Y
Tc = Tr
5 15 25 35 45
Time
Time

**Supplemental Movies**

**Movie S1. SOS and RAF in cells stimulated with EGF.**

**Movie S2. SOS and RAF translocation to the basal cell membrane in cells stimulated with EGF.** Time-lapse movies with a 1-min interval for 63 min. These two movies were acquired in the same field of view under epi (Movie S1) or total internal reflection (Movie S2) illumination. The left and right movies show signals from Halo (TMR)-SOS and GFP-RAF, respectively. Just before the 4th frame, 100 ng/ml (final concentration) of EGF was added to the observation medium. The field of view is 222 x 222 $\mu m^2$.
